# Genomic and epidemiological evidence for the emergence of a putative *L. donovani/L. infantum* hybrid with unusual epidemiology in Northern Italy

**DOI:** 10.1101/2023.08.09.552585

**Authors:** F. Bruno, G. Castelli, B. Li, S. Reale, E. Carra, F. Vitale, S. Scibetta, M. Calzolari, S. Varani, M. Ortalli, E. Franceschini, W. Gennari, G. Rugna, G.F. Späth

## Abstract

*Leishmania (L.) infantum* is the main causative agent of animal and human leishmaniasis in the Mediterranean basin. Despite its clinical significance, little is known on the genetic diversity of *L. infantum* parasites circulating in Italy. Here, we apply a comparative genomics approach on seven *L. infantum* isolates from different hosts (human, dog, cat, marten) and geographic regions (Emilia-Romagna, Sicily, Sardinia) as a first attempt to explore the breadth of parasite genetic heterogeneity in Italy. We revealed important genome instability at karyotype levels, with each isolate presenting a unique aneuploidy profile. Read depth analysis further identified strain-specific changes in gene dosage, which affected important virulence factors most of which are encoded by multi-copy gene arrays, such as amastins or surface antigen-like proteins. SNP-based clustering analysis of these genomes together with over 80 publicly available *L. infantum* and *L. donovani* genomes placed the Italian isolates into three geographically distinct clusters, with two isolates grouping with Spanish strains, two isolates grouping with Tunisian strains, and three isolates clustering with putative *L. infantum/L. donovani* hybrids isolated in Cyprus. As judged by microsatellite profiling of 73 isolates from dogs, sand flies and VL cases, these hybrid isolates are representative of a sub-population of parasites circulating in northeastern Italy that preferentially infects humans, but not dogs. In conclusion, our data uncover a remarkable heterogeneity of *L. infantum* isolates that indicates different geographic origin, including a novel hybrid-like genotype associated with an unusual infection pattern, placing Italy at the crossroad of *Leishmania* infection in the Mediterranean region.

## Introduction

Leishmaniases are a group of sand fly transmitted parasitic diseases that can affect mammals, including humans. In humans, these diseases are characterized by cutaneous, muco-cutaneous, or visceral tissue damage, the latter often being fatal if left untreated [1]. Pathogenesis and clinical outcome of *Leishmania* infection relies on a series of parasite-specific traits that ensure proliferation and fitness gain in different host systems.

First, these parasites have evolved different life cycle stages, including motile promastigote forms that are adapted for survival inside the midgut of phlebotomine sand flies, and intracellular amastigotes that infect immune cells of the reticuloendothelial system inside the mammalian host. Second, aside stage differentiation, *Leishmania* can further adapt to environmental variations encountered inside both insect and vertebrate hosts; this phenomenon has been linked to the intrinsic plasticity of the *Leishmania* genome and the genetic recombination that can follow the formation of parasite hybrids between strains and species. In the wake of next generation sequencing and the development of powerful computational pipelines for sequencing data analysis [2], comparative genomics approaches have provided important insight into the epidemiology of *Leishmania* infection in the field or parasite fitness gain in experiential settings. For example, comparative genomics has informed on (i) mechanisms underlying drug resistance and parasite evolution in treated patients [3–5], (ii) the origin, population structure, and phylogenetic relationship of clinical isolates [6–9], (iii) hybridization events and their effect on tissue tropism or clinical outcome of infection [10,11], (iv) the role of dynamic gene dosage changes in parasite fitness gain [12–15], or (v) the genomics of geographic adaptation [16,17], which is the scope of the current study.

*L. infantum* causes leishmaniasis in the Mediterranean Europe [18], with significant emergence of human cases observed in localized areas of Spain [19] and Italy [20–22]. Italy is hypo-endemic for human leishmaniasis, with less than one hundred cases of cutaneous (CL) and visceral leishmaniasis (VL) reported to WHO in 2020 [23], while dogs are considered the main reservoir hosts. Historically, leishmaniasis in Italy was restricted to the Tyrrhenian littoral, the southern peninsula, and the islands. However, in the last 20 years, sand fly vectors as well as human and canine *Leishmania* infections have been detected in northern Italy, traditionally classified as a cold area unsuitable for sand fly survival [24,25]. A recent study evaluated the spatio-temporal patterns of VL in Italy using records of the national Hospital Discharge Registers for 2009-2016 and indicated that VL is now endemic in the whole Italian peninsula, exhibiting a gradient of increasing incidence from north to south, peaking in Sicily [26,27].

In the last decade, an upsurge of human leishmaniasis cases has been observed in the region of Emilia-Romagna (RER), northeastern Italy [20–22]. However, there was no concurrent increase in the prevalence of canine leishmaniasis in the same period [28]. In addition, molecular typing studies in RER showed that the *Leishmania* strains circulating in dogs belonged to a different population compared to strains isolated from human VL cases [29,30] and sand flies [31]. Furthermore, the biting preference of *Phlebotomus (Ph) perfiliewi*, the suspected vector responsible for the increase of VL cases in RER, suggested the presence of a peri-urban or sylvatic reservoir other than dogs, able to spread the infection in such areas [32]. Consistently with this observation, a high frequency of *Leishmania* infection was recently identified in other potential reservoir hosts, including roe deer, hares, wolves, or red foxes [33].

While some studies have evaluated the epidemiology of VL in Italy, genetic characterization of *L. infantum* across different regions is lacking. Here, we combined comparative genomics and microsatellite profiling approaches on *L. infantum* field isolates that were obtained in Italy to shed first light onto parasite genetic heterogeneity. We provide evidence for a remarkable genetic diversity of *L. infantum* across the Italian peninsula and identify the strains from RER as putative *L. infantum/L. donovani* hybrids, which are closely related to Cypriote strains, thus correlating their unusual epidemiology and transmission pattern to their unique genetic constitution.

## Material and methods

### Clinical samples

Ten isolates of *L. infantum* were obtained from the cryobank of the National Reference Laboratory for Leishmaniasis (C.Re.Na.L., Palermo, Sicily). The strains were originally obtained from different hosts and geographies, including (i) from two naturally infected dogs by lymph node aspiration (MCAN/IT/2021/51327, Sardinia, referred to as leish3; MCAN/IT/2020/1265, Sicily, referred to as leish4), (ii) from the spleen of a wild marten (MMST/IT/2006/V2921, Sicily, referred to as leish5), (iii) from four owned cats by lymph node aspiration (MFEL/IT/2018/10816, Sicily, referred to as leish16; MFEL/IT/2018/CRENAL791; MFEL/IT/2018/CRENALBORIS; MFEL/IT/2018/CRENAL2147), and (iv) from bone marrow aspirates of three human VL cases from RER (MHOM/IT/2014/IZSLER-MO22, referred to as leish7; MHOM/IT/2014/IZSLER-MO23, referred to as leishMO23; MHOM/IT/2016/IZSLER-MO38, referred to as leishMO38). The strains leish3, leish4, leish5, leish7, leish16, leishMO23 and leishMO38 were used for whole genome sequencing analysis as described in Table 1.

**Table 1:**
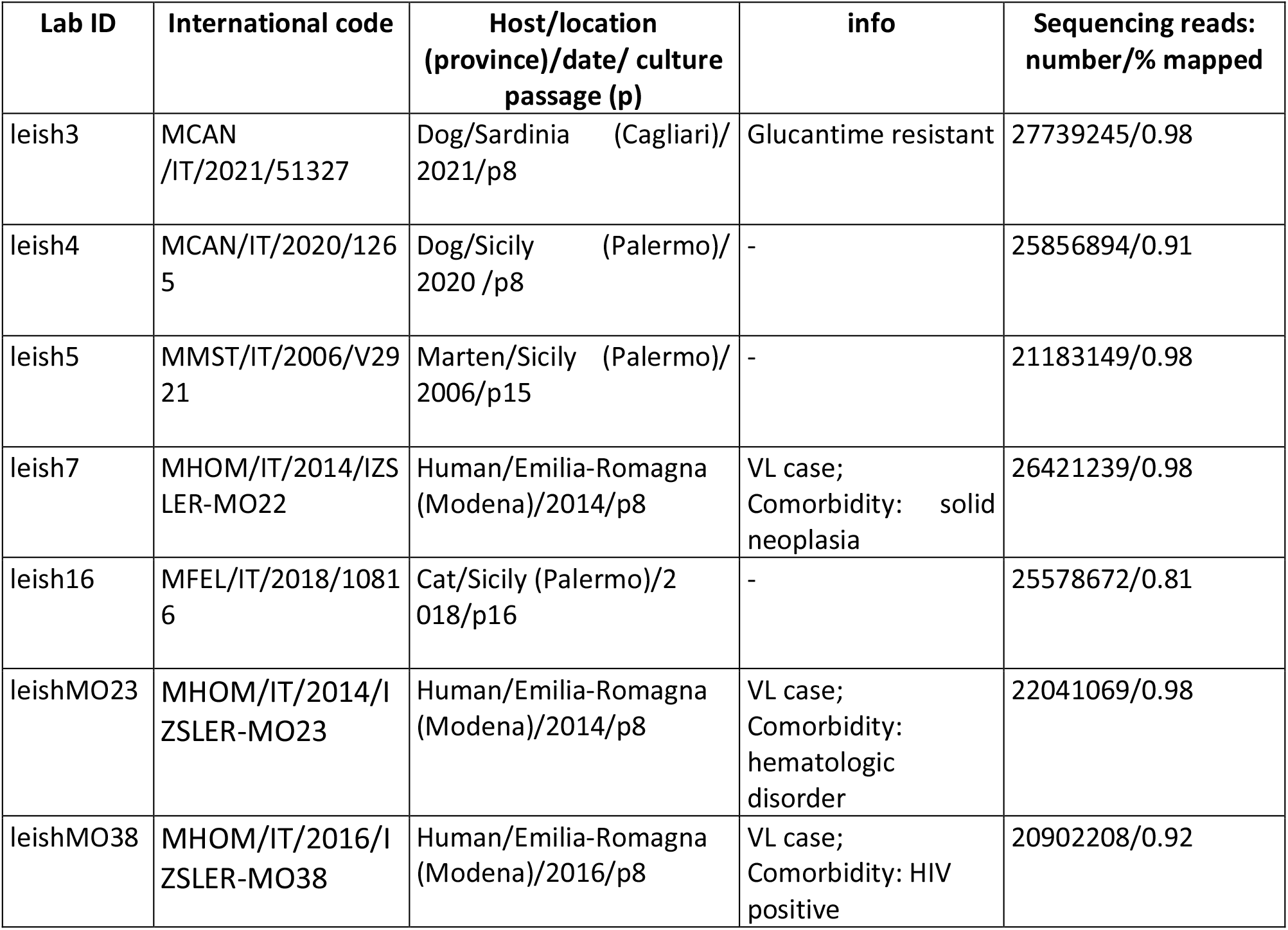
Overview of the *L. infantum* isolates that were analyzed in this study by whole genome sequencing.

### Parasite culture and DNA extraction

Parasites were grown in Evans’ modified Tobie’s medium (EMTM) supplemented with 10% fetal bovine serum, 5% sterile human urine, 1% phenol red solution, and 2% antibiotic solution (250 mg/ml gentamicin, and 500 mg/ml 5-fluorocytosine [34] At day seven, parasite viability and growth were assessed microscopically, and cells were used to inoculate 10 ml of RPMI-PY medium[34]. After reaching exponential growth phase (approx. 5 days), cells were pelleted by centrifugation at 2100 g for 10 min at 4 °C, and washed three times with physiological solution (0.9% NaCl) prior to DNA extraction according to a modified protocol described in Castelli *et al.* (2020) [35]. Briefly, cells were lysed by heat (96°C for 20 minutes) with 400 μl of a mixture containing 20% Chelex resin (Sigma, St. Louis, MO, USA), 1% Nonidet P-40 (Sigma, St. Louis, MO, USA), 1% Tween 20 (Sigma, St. Louis, MO, USA), and sterile distilled water. The mixture was centrifuged at 14,000 g for 10 minutes at 4°C and the DNA-containing, upper phase was collected and stored at - 20°C [36]. For multilocus microsatellite typing, DNA from sand fly samples was extracted using Qiagen DNeasy Blood & Tissue Kit (Qiagen, Hilden, Germany) according to the manufacturer’s instructions, and eluted in a final volume of 100 μl of elution buffer. DNA of strains isolated from VL cases was extracted from 100 μl whole blood (peripheral blood) or 100 μl bone marrow aspirate using NucliSENSeasyMAG (bioMerieux, Marcy l’Etoile, France).

### Genome sequencing

Whole genome DNA library preparation was performed using 100 ng DNA as input according to the standard protocol for Nextera DNA Flex Library Prep (Illumina, CA, USA) using Nextera DNA CD Indexes (Reference Guide 1000000025416.v07, Illumina). Quality and fragment size of all libraries were assessed by capillary electrophoresis on a TapeStation 4200, using a High Sensitivity DNA kit (Agilent, CA, USA). Individual DNA libraries were quantified using Qubit 2.0 Fluorometer with High Sensitivity dsDNA Assay (Thermo Fisher Scientific, MA, USA), and diluted to 4 nM with Resuspension Buffer. Normalized libraries were pooled using 10 μl per sample prior to sequencing, then the pooled library was diluted to a final concentration of 12 pM and denatured according to the sequencing protocol for V3 cartridges used with the Illumina MiSeq platform (Reference Guide 15039740.v10, Illumina). Samples were spiked with 1% of PhiX Control library and sequencing was run using a paired-end 300-cycle protocol.

### Computational analyses

WGS reads were mapped and sequencing data processed using GIP version 1.1.0 [2] (see Table S1 for mapping metrics). The outputs of the GIP pipeline were used for downstream analyses, ignoring the reads mapping to mitochondrial DNA (i.e. maxi and minicircles). Mean coverage in 300 bp bins as generated by the GIP pipeline were used to compute somy scores per chromosome by first normalizing bin scores for a sample by their median across the entire genome (to obtain comparable values between samples), multiplying by two (to scale somy values to the default diploid state assumed for most of the chromosome). These somy scores were represented on ‘per chromosome and per sample’ boxplots. The median somy score across bins belonging to a given chromosome were visualized on a sample versus chromosome heatmap.

Normalized mean coverages per gene are reported by GIP, as the mean coverage of the gene divided by the median coverage of the chromosome containing the gene. These values were plotted against gene indices, the index of a gene corresponding to its rank when sorted by genomic coordinates. These values were also used to compute a heatmap. In order to preserve a readable colour scale, genes having a normalized mean coverage below 0.5 or above 1.5 in at least one of the samples were not included in the heatmap.

SNP frequencies were retrieved from the filtered output of GIP for each sample. For a given set of samples, the union of all SNPs across samples was considered, assuming an alt-allele frequency of 0 when a SNP was missing from a given sample. Distributions of alt-allele frequencies were represented using violin plots, and pairwise scatterplots. Upset plots [37] were generated using the upsetplot 0.6.0 Python library, considering SNPs at alt-allele frequency 0 as missing from a given sample. SNPs were sorted according to genomic coordinates and assigned a corresponding index against which alt allele frequencies were plotted. Euclidean distances between samples in the SNP allele frequencies space were computed and used to build a dendrogram using the scipy.cluster.hierarchy module from the Scipy 1.8.0 Python library [38]. The clustering was obtained using the UPGMA method. Besides the more specialized Python libraries indicated above, post-GIP analyses were carried out using the Pandas 1.4.2 [39,40]; Matplotlib 3.5.1 [41] and Seaborn 0.11.2 [42] Python libraries.

### Quantitative real time (RT) PCR analysis

Copy number quantification for chr 8 was assessed by qPCR analysis using the SYBR® Green assay for the samples leish16 and leish3 and additional feline *Leishmania* strains (see above). The approach was based on a comparative quantitative detection between a first target, the histone deacetylase (HD) gene on chr 8, and a second target on diploid chr 28 used as control, corresponding to a gene encoding for a putative glycerophosphoryl diester phosphodiesterase (GDP). Briefly, total genomic DNA was extracted from 1×10^6^ parasites using DNeasy Blood & Tissue kit (Qiagen) following the manufacturer’s protocol. RT-PCR was performed in triplicates and carried out in 20 µl of reaction mixture containing 10 µl of Fast SYBR® Green Master Mix (Biorad), 0.2 µl (10 pmol/µl) of gene-specific forward and reverse primers (HD-2F TGTAGCAGCATGCGCGCGT and HD-2R CTAGACGCCGGGGCAATGA, generating a product of 475bp; GDP-2F GAGCACTTCACTCAA GAGGC and GDP-2R TCGTCGCTCTGTCAGCTGG, generating a product of 466bp), 2 µl of genomic DNA template and 7.6 µl nuclease free water to adjust the reaction volume. Negative controls (NTC) were included in all assays corresponding to reaction mixture without genomic DNA. Real-Time PCR was carried out in QuantStudio™ 3 Real-Time PCR System, 96-well (Life Technologies, Carlsbad, USA). The following thermal profile was used: pretreatment at 98 °C for 2 min then 40 cycles of 98 °C for 5 sec, 60 °C for 5 sec and a melting curve analysis to verify the amplification product.

### Microsatellite analysis of *Leishmania* samples from RER

The three *L. infantum* isolates from RER (leish7, leishMO23, leishMO38, see above) have been previously characterized by multilocus microsatellite typing (MLMT) using a panel of 15 dinucleotide microsatellite markers (Li41-56, Li46-67, Li21-34, Li22-35, Li23-41, Lm2TG, Lm4TA, Li71-5/2, LIST7039, Li71-33, Li71-7, CS20, Li45-24, TubCA and LIST7031) [30]. They were representative of a distinct genetic population of *L. infantum* circulating in humans and sand flies but not in dogs, defined as Population (Pop) B in [30]. In the present study, we expanded the epidemiologic characterization of the *L. infantum* strains circulating in RER by comparing their MLMT profiles to those of 30 *L. infantum* strains from VL (n=22) and sand flies (N=8) (Table S6) and 40 strains from dogs [30], for a total of 73 strains obtained from various areas of RER between 2013 and 2022.

### Population structure and phylogenetic analysis

MLMT profile results were analysed using models based on Bayesian clustering algorithm and genetic distances. Clustering analysis was performed by STRUCTURE v.2.3.4 software package [43], which determines genetically distinct populations on the basis of allele frequencies and estimates the individual membership coefficient (Q-value) in each probabilistic population. The Markov chain Monte Carlo iterations were set to 200,000 and the length of burn-in period to 20,000. For each value of K (estimated number of populations) between K=1 and K=10, ten independent simulations were performed. To estimate the best number of populations, the likelihood values computed by STRUCTURE were analyzed by STRUCTURE HARVESTER v0.6.1 [44] implementing the Evanno’s ΔK statistic, which is based on the rate of change in the log probability of data between successive K values [45]. Data were sorted by CLUMPP 1.1.2 software [46] and the barplots of the CLUMPP outfiles were visualized using an online tool, STRUCTURE PLOT [47]. Descriptive statistics for genetic populations were calculated by the GDA software [48]; this included mean number of alleles (MNA) per population, proportion of polymorphic loci (P), expected (He) and observed (Ho) heterozygosity and inbreeding coefficient (Fis). Phylogenetic analysis was performed based on microsatellite genetic distances using the application BEASTvntr package implemented in the BEAST2. 2.4.3 [49]. The diploid data were entered as two distinct partitions, with a linked tree and strict clock; the Sainudiin mutation model of microsatellites [50] was then selected with a chain length of ten million steps. Trees were visualized with iTOL v6 (https://itol.embl.de/).

A phylogenetic tree based on 14 coincident markers was created as described above for a global microsatellite analysis, and it included the MLMT profiles of the 73 strains from ER and those of 65 strains of the *L. donovani* complex, including (i) 11 strains from other endemic Italian regions [30], (ii) 51 strains from other countries (Table S7) [51–55], and (iii) 3 WHO *L. infantum* reference strains, namely MHOM/TN/80/IPT1 (MON-1), MHOM/IT/86/ISS218 (MON-72), and MHOM/DZ/82/LIPA59 (MON-24) (Table S6), which represent the most common zymodemes responsible for leishmaniasis in Italy [56].

## Results

### Cultured *L. infantum* isolates show strain-specific and convergent aneuploidies

We applied a comparative genomics approach to gain first insight into the genetic diversity of *L. infantum* field isolates in Italy. We deliberately chose initially five isolates from different hosts (human, dog, cat, marten) and geographic regions (RER in north-eastern Italy, Sicily, Sardinia) to maximize the assessment of parasite genetic heterogeneity (Table 1). Following culture expansion, DNA extraction and Illumina short read sequencing, we applied our genome instability pipeline (GIP, [2]) to map the reads on the JPCM5 *L. infantum* reference genome [57] and assess read depth variations for each individual sample. We then compared chromosome read depth variations between samples using per chromosome boxplots of normalized coverage (somy score) in 300 bp genomic bins (see methods). As judged by changes in the somy score, all samples showed important karyotypic variations thus confirming intrinsic instability of the *L. infantum* genome as previously reported [58,59] (Figure 1A). As expected from previous studies, all samples were tetrasomic for chromosome (chr) 31 [60] while displaying unique karyotypic profiles considering the other chromosomes. This diversity may be largely attributed to strain-specific selection of different aneuploidies during culture adaptation, which we previously linked to *in vitro* fitness gain [12–15]. The heat map shown in Figure 1B reveals leish4 as the karyotypically most stable sample, with most chromosomes showing a median somy score between 2 to 2.5. All other samples show more important somy variations, including full trisomies or higher, mosaic aneuplodies. Surprisingly, the propensity of amplification seems chromosome-specific: several chromosomes maintain a disomic state in all cultured samples (chr 19, 21, 25, 27, 28, 30, 32), while others show significant increase in somy score across most or all samples (e.g. chr 5, 9, 20, 26), indicating convergent selection of these aneuploidies.

**Figure 1:**
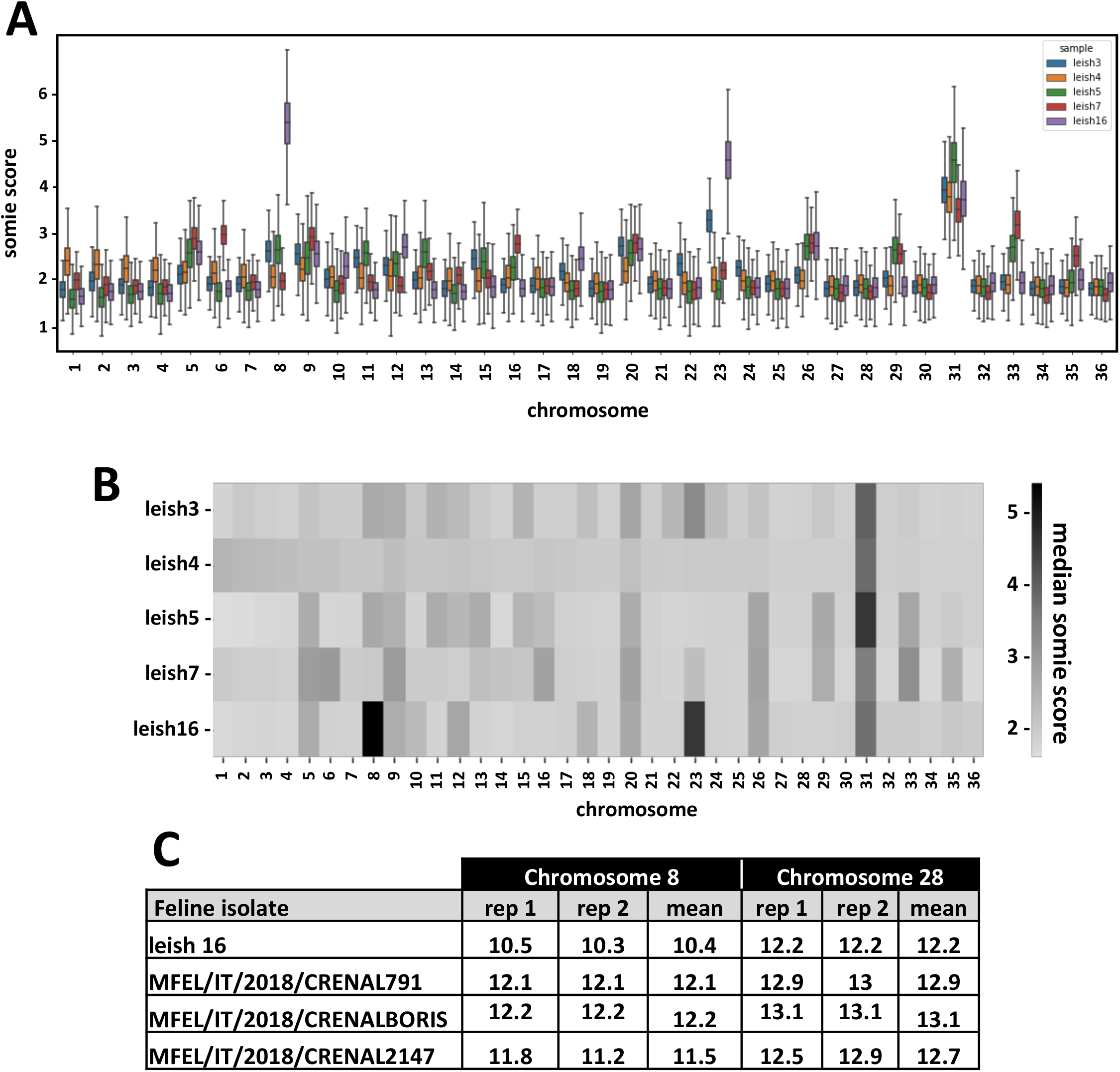
Karyotypic analyses of the *L. infantum* isolates (n=5). (A) Box plot representing the normalized sequencing coverage distributions for each chromosome (X-axis) expressed as somy score (Y-axis). The horizontal line in each box indicates the median normalized coverage value. The lower and upper edges of the box show respectively the lower quartile and upper quartile of normalized coverage values. The whiskers show maximum and minimum coverage values excluding outliers, which are not shown to ease readability. Different strains are shown in different colors (see legend in the figure). (B) Heatmap showing the median somy score for the different strains across the 36 chromosomes. Darker tonalities reflect higher somy values. (C) qPCR analysis with SYBR® Green assay to quantify chr 8 copy number in feline *Leishmania* strains. CT values are shown for the histone deacetylase (HD) gene on chr 8 and the glycerophosphoryl diester phosphodiesterase (GDP) gene on disomic chr 28 used as a control.

Our data revealed pentasomies for chr 8 and 23 in the leish16 sample. Given that this sample represents the first *L. infantum* genome sequenced from a feline isolate, we investigated if this unusual amplification pattern may be selected in this particular host. This possibility was ruled out by quantification of chr 8 copy number by qPCR analysis of three independent feline isolates, which showed a disomic amplification score comparable to disomic chr 28 included as a control (Figure 1C). Thus, the pentasomy for chr 8 may either have been uniquely selected in leish16 during feline infection or underwent important amplification during the 16 culture passages of this sample (see Table 1).

Thus, the Italian *L. infantum* field isolates show important somy variations in culture compared to the haploid reference genome. Convergent amplification of a small number of chromosomes in most or all strains suggests that these aneuploidies are a potential driver of *L. infantum* fitness gain in culture. Surprisingly, similar chromosomes were linked to culture adaptation of *L. donovani* isolates from the Indian sub-continent and from Sudan (e.g. chr 5, 20, 26) [12], suggesting conserved fitness mechanisms between both parasite species when selected for growth in culture. Aside these convergent aneuploidies, we further observed strain-specific, karyotypic changes that may fine-tune adaptation in response to other genetic differences between the strains, which we further analysed in the following.

### Gene copy number variation (CNVs) in the *L. infantum* isolates affect known *Leishmania* virulence genes

We next examined the gene-level coverage information generated by GIP to assess strain-specific and convergent gene dosage changes with respect to the JPCM5 reference genome (Table S2). Unlike karyotypic changes that are rapidly selected during short-term culture adaptation (5 to 20 passages), changes in gene CNVs are much less dynamic and thus can reveal gene dosage changes that were selected in the field. Plotting the normalized read depth coverage per gene, we observed a wavy pattern across the genomes for all isolates, with lower read density observed at the chromosome telomers and a single peak in read depth density observed towards the center of each chromosome (Figure 2A and Figure S1). The most parsimonious explanation for this pattern involves (i) depletion of reads with low MAPQ scores due to non-unique mapping on telomeric repeats, and (ii) augmentation of reads at the origins of replication as a result of active DNA synthesis in growing parasites [61].

**Figure 2:**
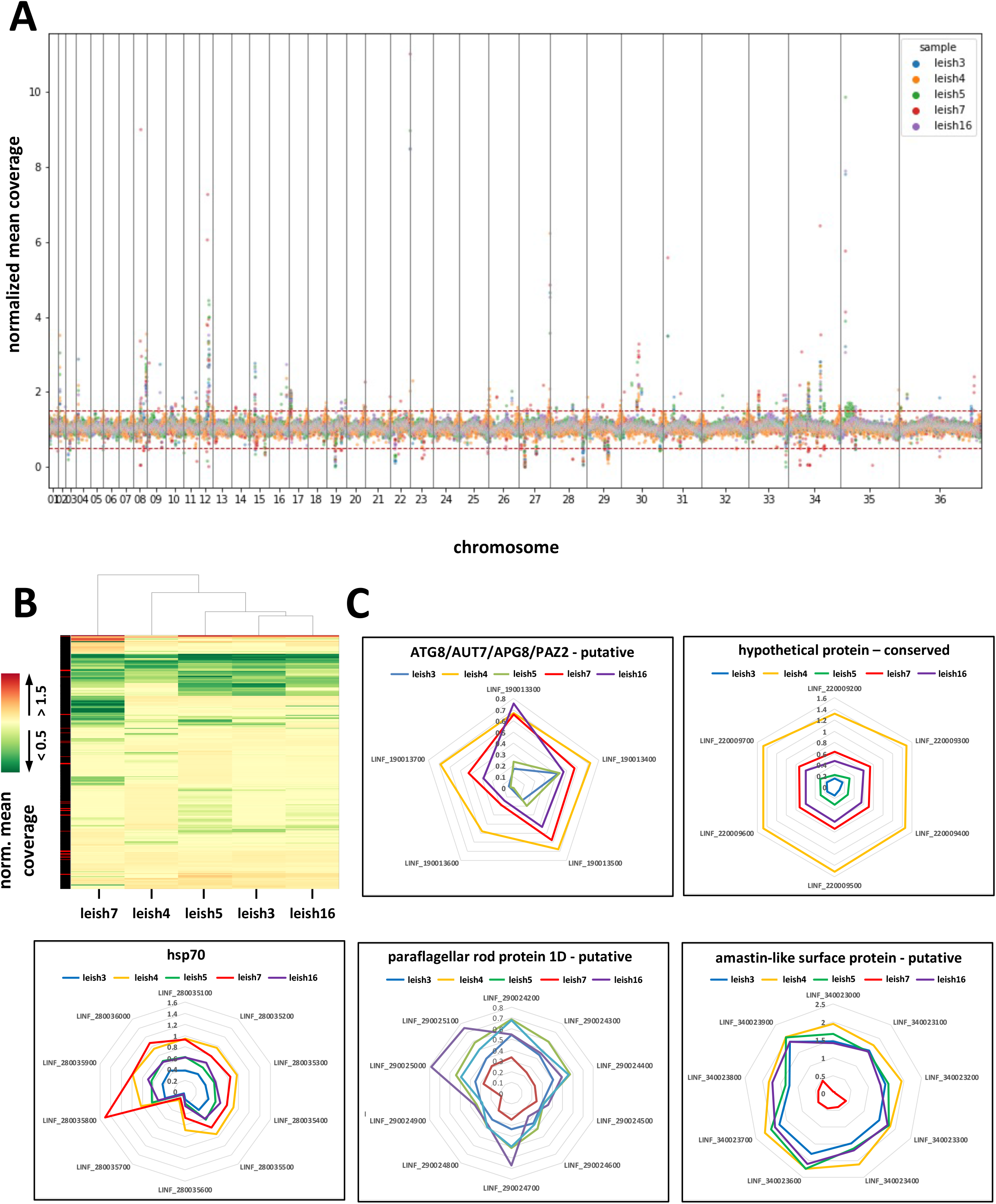
Gene copy number analyses of the *L. infantum* isolates (n=5). (A) Scatterplot showing the normalised mean coverage (Y-axis) for each gene across the 36 chromosomes (X-axis). Different strains are shown in different colors (see legend in the figure). (B) Heatmap of normalised mean coverage. Gene depletions and deletions are indicated in green and gene amplifications in red. The column on the left indicates the MAPQ score (red, MABQ<40; black, MAPQ>40). (C) Radar plots of the normalised mean coverage values for the genes indicated. Different strains are shown in different colors (see legend in the figure).

Second, we observed a series of gene CNVs that correspond to amplification or depletion of gene copies with respect to the reference genome, but also in-between the 5 samples. Significantly, these variations were not scattered across the genome, but localized in defined genomic regions that were similar in all samples, thus revealing possible hot spots of gene dosage changes (Figure 2A). Closer inspection of the 50 most significant gene CNVs identified both single gene and sub-chromosmal amplifications, many of which have been previously linked to infectivity, including amastins [62], surface antigen-like proteins [63–65] or genes implicated in the synthesis of phosphoglycan virulence factors [66–68], such as phosphoglycan beta 1-3 galactosyltransferase genes (LINF_020006900, LINF_020007000) or ppg3 (LINF_350010100, LINF_350010200) encoding filamentous proteophosphoglycan [69] (Figure S2A, Table S3). Significantly, most gene amplifications are observed across several isolates, suggesting convergent evolution by natural selection. Convergence is not only observed at the gene level but also the level of gene function, with amplified genes often sharing the same annotation even though they are on different chromosomes and encoding different proteins as shown for example for the genes encoding surface antigen proteins and amastins (Figure S2B).

We also identified 116 gene depletions (normalized mean coverage < 0.5) and deletions (normalized mean coverage = 0) that were shared or strain-specific in the five *L. infantum* isolates (Figure 2B). Most of these changes occurred in multi-copy gene arrays (Table S4), where recombination events between identical sequences allow for dynamic gene copy number changes. In particular, such dynamic variations between the strains were observed for (i) five gene copies on chr 9 encoding for the autophagy gene ATG8, which has been implicated in *Leishmania* stress response and infectivity [70], (ii) six gene copies on chr 22 encoding for an uncharacterized, hypothetical protein, (iii) a cluster of nine amastin genes encoded on chr 34, known to control amastigote infectivity [62], (iv) 10 HSP70 gene copies encoded on chr 28 that we previously revealed as potential hot spot of environment-genotype interaction in *L. donovani* field isolates [71] and (v) 10 genes on chr 34 encoding for paraflagellar rod protein (Figure 2C).

The reduced read depth that we observed for individual gene copies suggests mosaic gene CNVs, i.e. mixed populations of wildtype and deleted genotypes. Only few genes show true deletions as judged by the absence of reads, including LINF_270010900 encoding for a hypothetical gene (deleted in leish16, leish3, leish5 and leish7), or the ATG8 gene LINF_190013600 and Flagellar Member 8 gene LINF_330040600 in leish5 (Figure 2C). The penetrance of these deletions within non-clonal, heterogenous parasite populations suggests selection against these genes, even though loss due to genetic drift cannot be ruled out.

### Discovery of leish7 as highly divergent *L. infantum* strain

We next assessed the evolutionary relationship between our five *L. infantum* isolates comparing SNP numbers, positions and frequencies from the GIP output. Based on the number of alternative (alt) alleles, leish16 is the most similar to the JPCM5 reference genome with only 402 SNPs identified, while leish7 is the most divergent with 44,916 SNPs (Figure 3A and B). Cluster analysis based on these variants revealed highest similarities between leish4 and leish16 on one hand, and leish3 and leish5 on the other hand, while leish7 represents its own branch, with similar distance from either of the two other clusters (Figure 3C). Plotting frequency against location revealed defined patches of heterozygous SNPs at a frequency of 50%, suggesting the presence of distinct haplotypes that may be remnants of former hybridization events.

**Figure 3:**
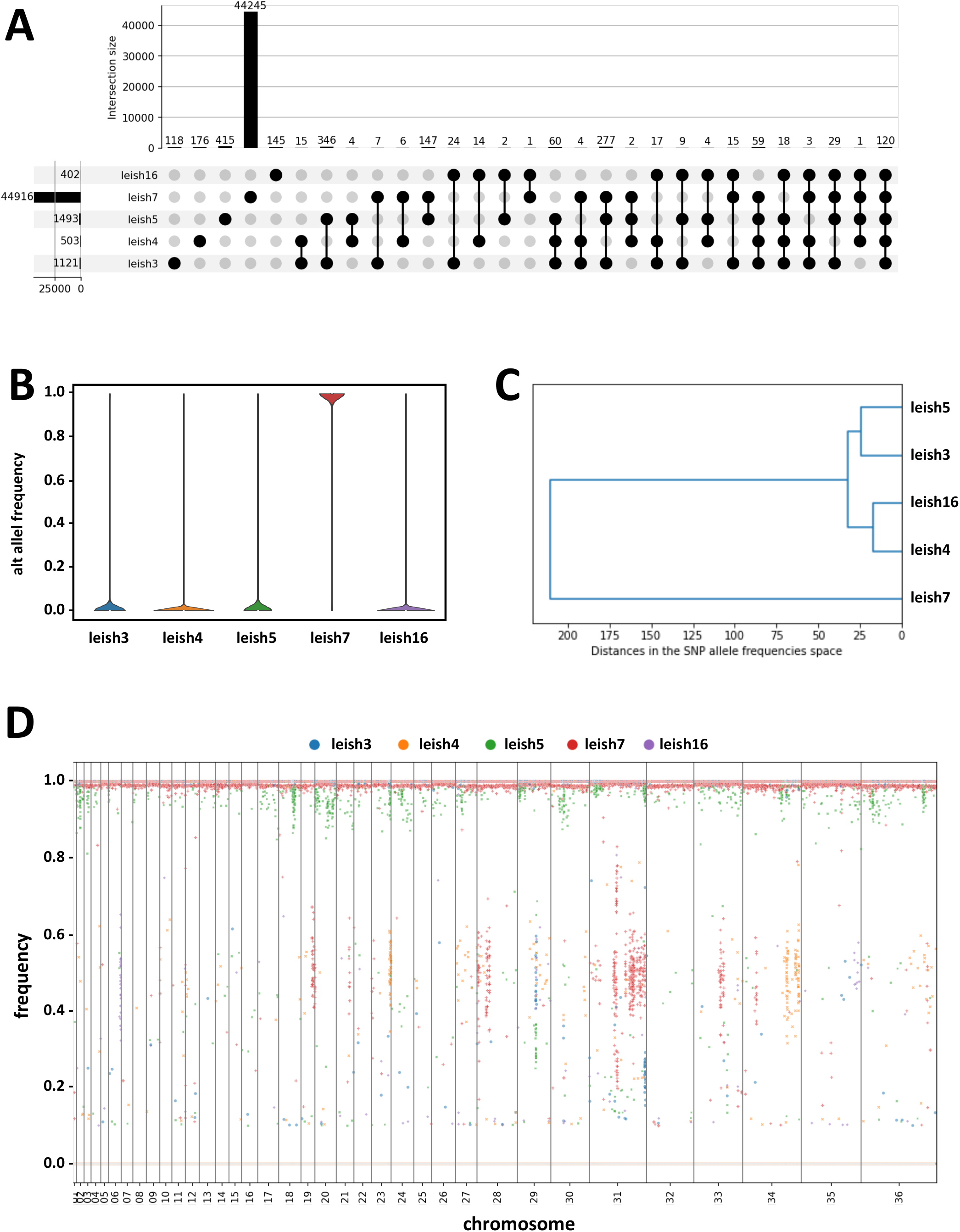
Genome-wide SNP analyses of the *L. infantum* isolates (n=5). (A) Upset plot of SNPs identified in the different strains. The bars visualize the number of SNPs common to a given combination of samples. The numbers in the bar plot indicate shared SNPs for the given combination. The numbers left to the strain ID indicate the total number of SNPs in each isolate. (B) Violin plot representing density estimates of the distribution of alt allele frequencies for the indicated samples. (C) Cluster analysis based on pairwise Euclidean distances between alt allele frequencies across SNPs. The tree is constructed from those distances using the UPGMA method. (D) The scatterplots display individual SNPs as dots. The x-axis indicates the genomic index (rank of the SNP in a genomic sort) while the y-axis indicates the variant allele frequency. The different chromosomes are displayed one after the other, and their boundaries are visualized as vertical lines. The individual SNPs are colored according to the samples as indicated by the legend in the graph.

### Investigating the possible origin of the leish7 putative hybrid

We next established the genome sequence for two additional isolates (leishMO23 and leishMO38) obtained from human VL cases living in RER, northeastern Italy. Cluster analyses revealed their close relationship to leish7, confirming transmission of a distinct parasite sub-population – likely a hybrid – in RER (Figure 4A, top right insert). We further expanded the cluster analysis including 11 genomes that we previously generated as part of the LeiSHield project (www.leishield.org, [58,59]) and 63 genomes analyzed by Franssen *et al.* (2020) [72]. While four of our Italian *L. infantum* strains clustered with isolates from Tunisia (leish3 and leish5) and Spain (leish4 and leish16), the three RER hybrid-like strains (leish7, leishMO23, leishMO38) clustered with the known *L. infantum/L. donovani* hybrids first described in Cyprus (Linf_CH33, 35, 36 and Ldo_CH33). Closer inspection of shared and unique SNPs between these hybrids (Figure 4B) established a clear common origin, with over 34,000 SNPs shared between the geographically distinct hybrid-like strains (out of 44,961 to 49,274 SNPs). Over 5,000 SNPs distinguished the RER strains from the Cypriote hybrids, indicating divergent evolution that may be driven by ecologic variations within the two distant locations.

**Figure 4.**
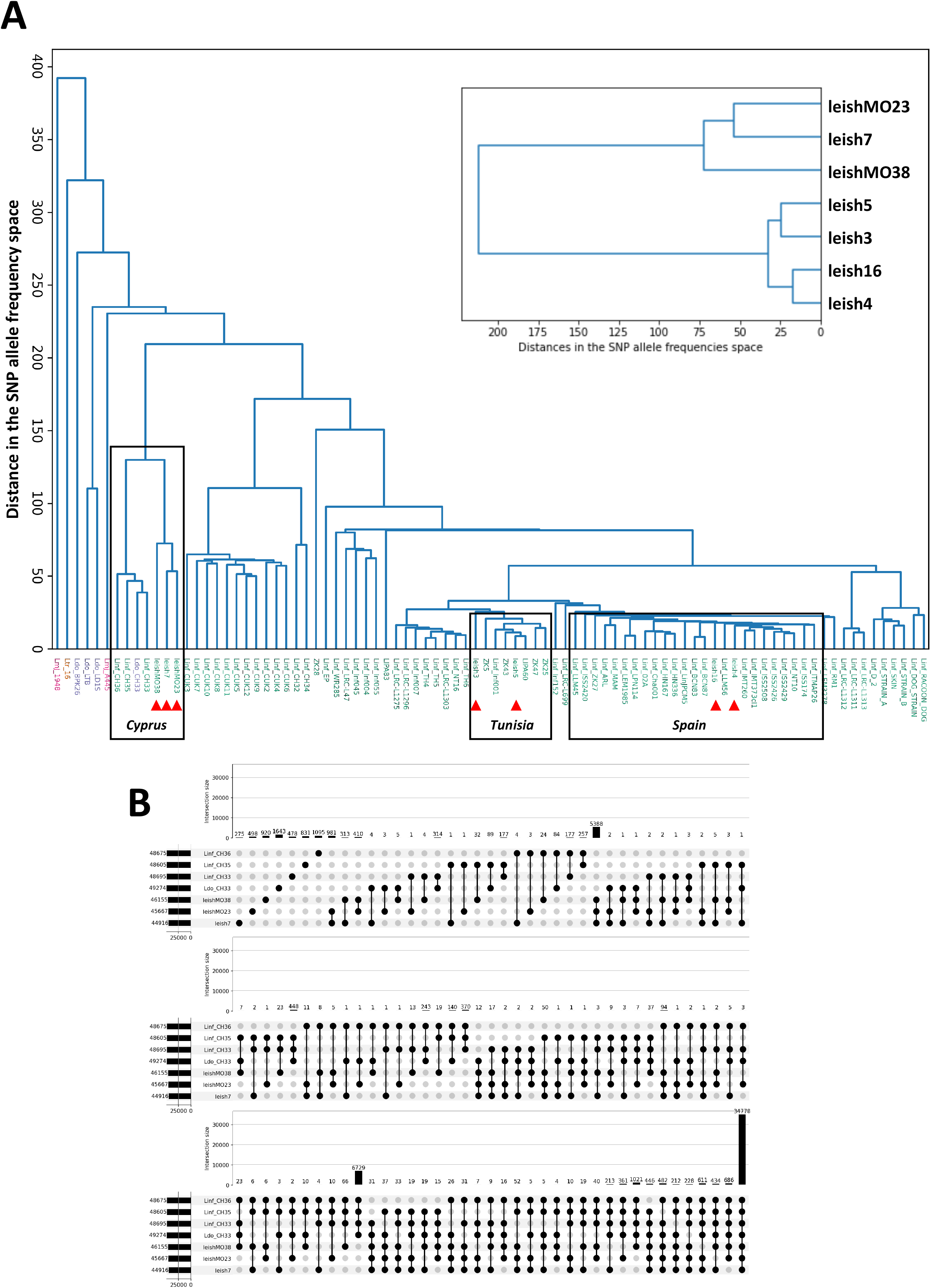
Putative origin of the leish7 hybrid based on SNP analysis. (A) Cluster analysis based on pairwise Euclidean distances between alt allele frequencies across SNPs. The tree is constructed from those distances using the UPGMA method. On the main tree, sample names are colored according to their recorded species. The samples that were not sequenced in the present study have their names prefixed by a tag indicating the species: Lmj (*major*), Ltr (*tropica*), Ldo (*donovani*) or Linf (*infantum*). The samples that were sequenced in the present study are indicated by a red triangle. The top right insert shows the cluster analysis of two additional hybrid strains (leishMO23 and leishMO38) in comparison to the other strains sequenced in this study. (B) Upset plot of SNPs identified in the different strains indicated. The bars visualize the number of SNPs common to a given combination of samples. The numbers in the bar plot indicate shared SNPs for the given combination. The numbers left to the strain ID indicate the total number of SNPs in a given isolate.

### Population structure and phylogenetic analysis of putative hybrid strains from northeastern Italy

In order to better define the prevalence of the leish7 hybrid-like genotype in notheastern Italy, microsatellite analysis was performed on 73 *L. infantum* strains including 22 human VL cases, 8 sand flies and 40 canine cases from various areas of RER. Bayesian clustering as implemented in STRUCTURE assigned the 73 MLMT profiles from these *L. infantum* strains to two main populations (the highest value of ΔK was for K=2; Fig. 5A), namely Population A (PopA) that consisted of canine *L. infantum* strains only, and Population B (PopB) that consisted of all the *L. infantum* strains from VL (including leish7, leishMO23 and leishMO38) and sand flies (Fig. 5B). Additionally, the microsatellite markers that were analyzed in the PopB showed low average expected heterozygosity (He=0.311) and observed heterozygosity (Ho=0.043), and a mean inbreeding coefficient of 0.864 (Table 2).

**Figure 5.**
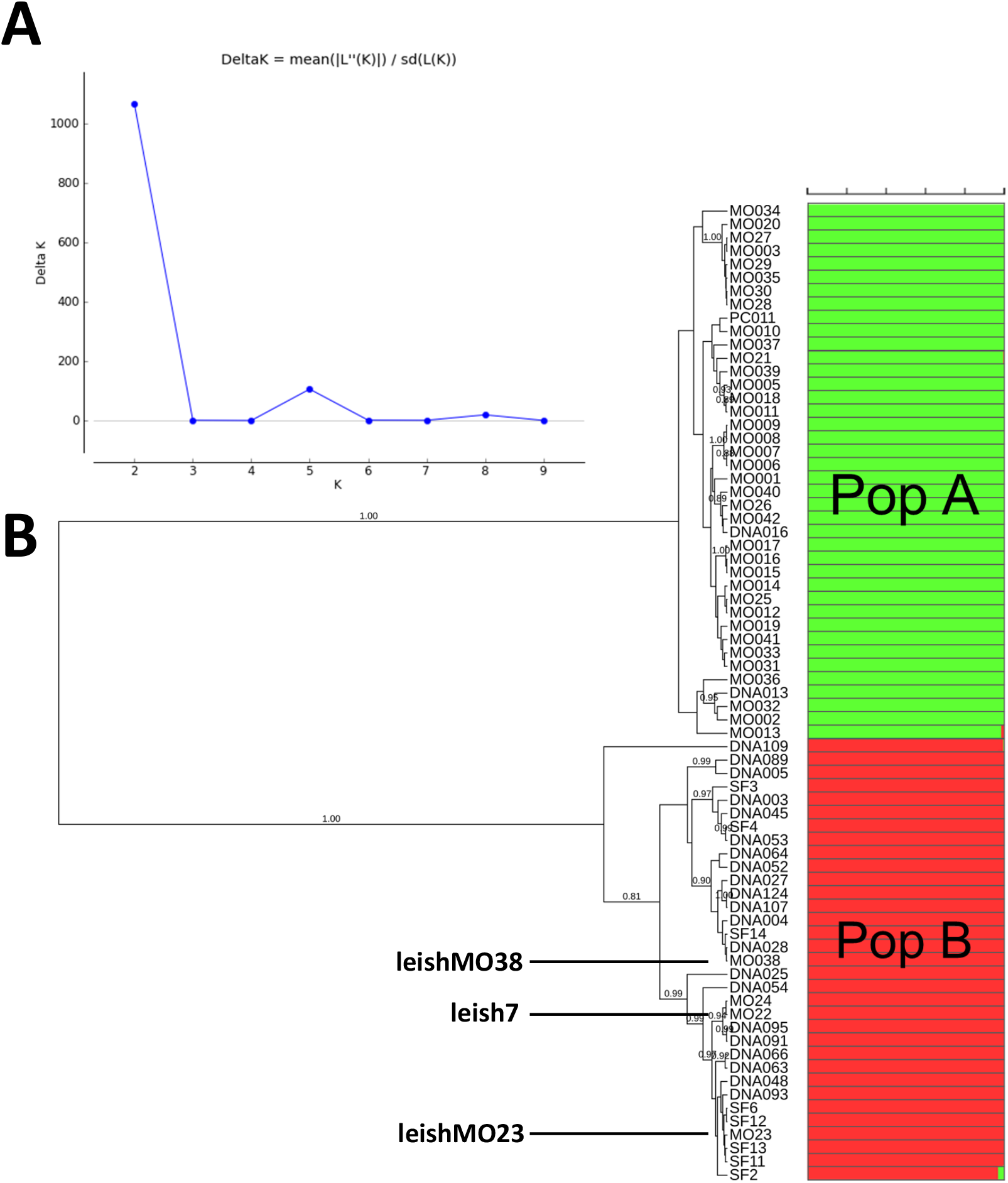
Combined population structure and genetic distances analyses using data of 15 microsatellite loci for the 73 *L. infantum* strains from the Emilia-Romagna region (northeastern Italy). (A) Plot of the ΔK values per populations (K) according to the Evanno method [45] on data generated by STRUCTURE analysis. The primary peak at K=2 indicates the existence of two populations in the investigated strains; (B) Bayesian phylogenetic tree obtained by the Sainudiin model [50]. Posterior probability values >0.80 are indicated at the nodes. Populations as inferred by STRUCTURE are indicated by colored bars: green for PopA, red for PopB. The isolates that were sequenced in this study (leish7, leishMO23 and leishMO38) are indicated in bold and bigger font size.

**Table 2.** Population genetic characterization of the *L. infantum* population B, including strains from VL cases and sand flies in the Emilia-Romagna region, northeastern Italy.

| Population | N | P | MNA | He | Ho | F<sub>is</sub> |
| --- | --- | --- | --- | --- | --- | --- |
| B | 33 | 0.867 | 2.600 | 0.311 | 0.043 | 0.864 |

Footnote: Population, cluster as inferred by STRUCTURE analysis; N, sample size; P, proportion of polymorphic loci; MNA, mean number of alleles; He, expected heterozygosity; Ho observed heterozygosity; F<sub>is</sub>, inbreeding coefficient

The global phylogenetic analysis showing the position of human and sand fly strains from RER within the *L. donovani* complex is represented in Figure 6. Canine *L. infantum* strains from RER belonging to the original PopA were grouped in a monophyletic clade together with VL strains from other Italian regions and Mediterranean strains belonging to *L. infantum* zymodemes MON-1, MON-72, and MON-98. On the contrary, *L. infantum* strains of VL and sand flies from RER belonging to PopB grouped in a monophyletic clade that was intermediate between *L. infantum* and *L. donovani* clusters and that included two strains from Cyprus, namely CH35 and CD44cl.1, both classified as *L. donovani* MON-37 [73], thus clearly establishing the origin and hybrid-like nature of the PopB parasite isolates.

**Figure 6.**
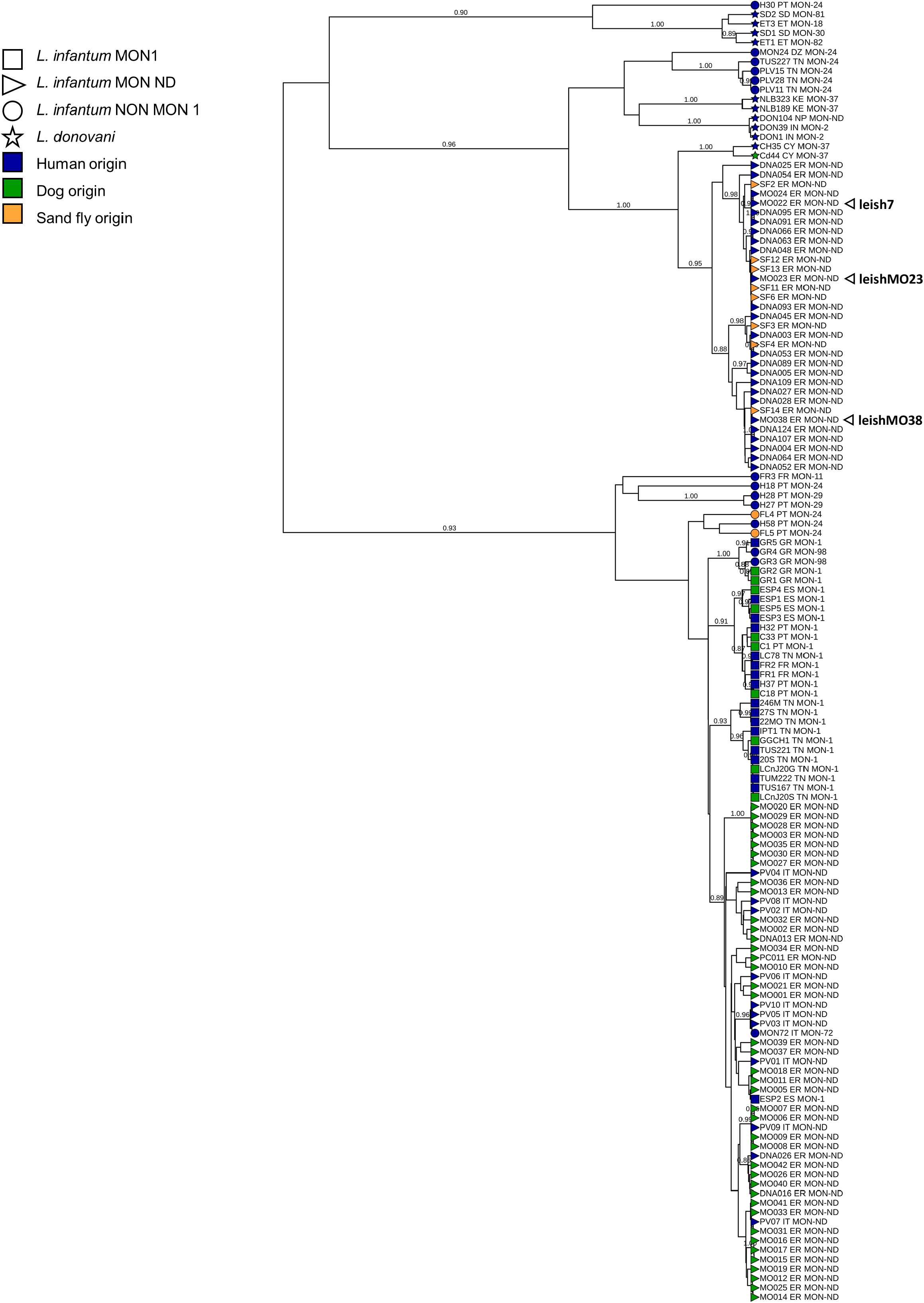
Bayesian phylogenetic tree of 73 L. infantum strains from the Emilia-Romagna region, 65 MLMT profiles available in the literature and 3 WHO reference strains. The tree was obtained by the Sainudiin model [50] using data of 14 coincident microsatellite loci. Posterior probabilities > 0.80 are showed near the nodes. Strains designations specify, respectively, the laboratory code, the zymodeme (MON1 indicates strains belonging to the MON-1 zymodeme, NON MON1 indicates strains that do not belong to the MON1 zymodeme, and MON ND indicates not defined strains), the geographical origin (IT, Italy; ER, Emilia-Romagna region; CY, Cyprus; FR, France; SP, Spain; GR, Greece; PT, Portugal; DZ, Algeria; TN, Tunisia; KE, Kenya; SD, Sudan; ET, Ethiopia; IN, India; NP, Nepal). The isolates that were sequenced in this study (leish7, leishMO23 and leishMO38) are indicated in bold and bigger font size.

## Discussion

The present study combines comparative genomics with molecular epidemiology approaches to deliver a first comprehensive, high-resolution investigation of intraspecific genetic diversity and heterozygosity within the *L. infantum* species in Italy, one of the most important *Leishmania* species worldwide from both a veterinary and public health perspective [27,74,75]. We revealed a remarkable heterogeneity in Italian *L. infantum* isolates that are related to various parasite genotypes of different geographic origin, including a hybrid-like genotype first described in isolates from Cyprus.

Our results place Italy at the crossroad of *L. infantum* infection in the Mediterranean and inform on the possible origin of regional sub-populations and their local transmission cycles. We identified a series of *L. infantum* strains from Sardinia and Sicily that are genetically close to those from Spain (south-western, SW Europe) and Tunisia (North Africa), highlighting Italy’s central location in the Mediterranean Basin and its historical and current role as one of the main routes of human migration. Based on genetic similarity, the strains leish3 and leish5 likely belong to the North African MON-1 population endemic for southern and eastern Mediterranean regions [53,76]. This population is genetically different from the European MON-1 population found in SW Europe and South America, which corresponds to the strains leish4 and leish16 and includes the JPCM5 strain used as a reference in this study [53,76].

The complex *Leishmania* epidemiology is the result of the parasite’s transmission dynamics, the distribution and geographic abundance of competent vectors, past and current exposure of human populations to the parasite, environmental parameters that impact the seasonal activity and distribution of sand flies (e.g. air temperature, relative humidity, ground cover, altitude, vegetation), and the presence of reservoir hosts [27,77–79]. In Italy, as in the whole Mediterranean basin, dogs (*Canis lupus familiaris*) are incriminated as the primary reservoir hosts of zoonotic human leishmaniasis [24], even if other domestic and wild mammals infected with *L. infantum* could play a possible role in the biological cycle of the parasite [80,81]. When compared to *L. donovani* species, *L. infantum* MON-1, the most common zymodeme in dogs and humans, has been considered a genomically rather homogeneous population regardless of geographic distribution [72]. The parasite’s adaptation to dogs may have been an evolutionary advantage, and the frequent mobility of the canine reservoir may have contributed to the spread of the *L. infantum* MON-1 group [82]. However, microsatellite analyses and WGS showed that the MON-1 “core” *L. infantum* clade displays a certain degree of isolation by distance. Three distinct populations have been shown to circulate within the Mediterranean Region, i.e. SW Europe, North Africa, and South-Eastern (SE) Europe [72,76]. Thus, the genetic structure within *L. infantum* MON-1 population is mainly related to the parasites’ geographic origin, echoing our previous epidemiological investigation conducted in Tunisia that revealed ecology rather than clinical phenotype as a key driver of parasite genomic adaptation [59].

Our investigation further uncovered putative hybrid strains from RER (leish7, leishMO23, and leishMO38), which may be linked to the emergence of VL in this region. Even though human and canine leishmaniasis cases have been reported in the past mostly from central and southern Italy, typically characterized by hot summers and mild winters [27,35,74,83], human leishmaniasis is recently emerging in the northern part of the country, including RER [29,84]. Our MLMT and whole genome sequencing analyses demonstrated the transmission in RER of putative hybrid populations that are closely related to divergent *L. infantum* strains first described in Cyprus as judged by the presence of 34,778 shared SNPs (74% of the total SNP count). Previous analyses described the Cypriote strains (e.g. strain CH33) as a novel *L. donovani* sensu lato (s.l.) group based on its unusual MON-37 zymodeme [73] [51,85,86], which may either represent a distinct evolutionary lineage within the *L. donovani* complex or an ancient hybrid between *L. infantum* and *L. donovani* [72]. The presence of over 11,000 SNPs that distinguish the RER and Cypriote hybrids indicates genetic differentiation since the split from their common ancestor. Indeed, the *L. infantum* hybrid population circulating in humans and sand flies in RER (PopB) is characterized by a high inbreeding coefficient and a low observed heterozygosity, suggesting intraspecific recombination or selfing, which likely contributed to the genetic distance observed today to the hybrid strains circulating in Cyprus.

Similar to Cyprus, leishmaniasis in RER displays a peculiar epidemiology. The first documented epidemic of VL dates back to 1971-1972 [87], two decades before the reported northward spread of canine leishmaniasis in Italy [24]. In addition to the extreme summer dryness of 1971, which likely contributed to an excess of sand flies and increased human exposure to their bites, the epidemiologic analysis led to the hypothesis that a cryptic infection in an unidentified domestic or wild animal reservoir was the cause of the outbreak [87]. Following a sharp decline in human leishmaniasis cases in the ensuing decades, RER has seen a re-emergence of VL cases since 2010 [20–22]. Currently, the area is experiencing two distinct epidemiological *Leishmania* transmission cycles, one of which primarily affects humans involving *Ph. perfiliewi* as a vector (PopB by MLMT analysis) and the other which affects dogs (PopA). A previous study showed no genetic flow between these two parasite populations [30] despite their geographic overlap, incriminating distinct transmission cycles that likely involve specific vector species and/or reservoir hosts [32,33].

In conclusion, our genomic and epidemiologic data uncover a remarkable heterogeneity of *L. infantum* isolates in Italy, including a hybrid genotype reminiscent to Cypriote strains. The unusual infection pattern of this putative hybrid highlights the public health risk of introducing new *Leishmania* species due to the increased flow of migrants and tourists from countries where non-European species are endemic [88–90]. The risk of active transmission of newly introduced parasite species may be limited by the absence of permissive reservoir hosts or the lack of competent, local *Phlebotomus* species. However, this scenario does not apply for parasite species associated with an anthroponotic cycle of transmission, such as *L. tropica* and *L. donovani* [88,91]. In addition, *Leishmania* has a remarkable adaptation capacity to highly diverse ecologies, as illustrated by the introduction of *L. infantum* from the Old World to the New World during the South American conquest, that led to important epidemiological consequences. [17,92]. The challenge of future studies lies in the application of a trans-disciplinary approach combining genomics, epidemiology, ecology and entomology investigations to identify distinct transmission cycles and the underlying environmental parameters that drive the evolution of distinct, region-specific *L. infantum* sub-populations with unique clinical features.

## Supporting information

Table S1: Mapping metrics

Table S2: All gene CNVs

Table S3: Most significant gene CNVs

Table S4: Gene depletions

Table S5: SNP frequencies

Table S6: Designation, characteristics and MLMT profiles of the Leishmania strains

Table S7: Leishmania donovani complex strains used in this study

## Aknowledgments

This research was funded by the EU H2020 project LeiSHield-MATI-REP-778298-1, the Ministry of Health, Italy (grant E54I19002870001—IZSLER PRC2019016, grant number “IZS SI RC SI 04/21” and GR-2019-12369134 “Leishmaniasis in Agrigento, Caltanissetta and Palermo provinces: human outbreaks and animal reservoirs”), by EU funding within the NextGeneration EU-MUR PNRR Extended Partnership initiative on Emerging Infectious Diseases (Project no. PE00000007, INF-ACT).

## Supplementary Figure legends

**Figure S1:**
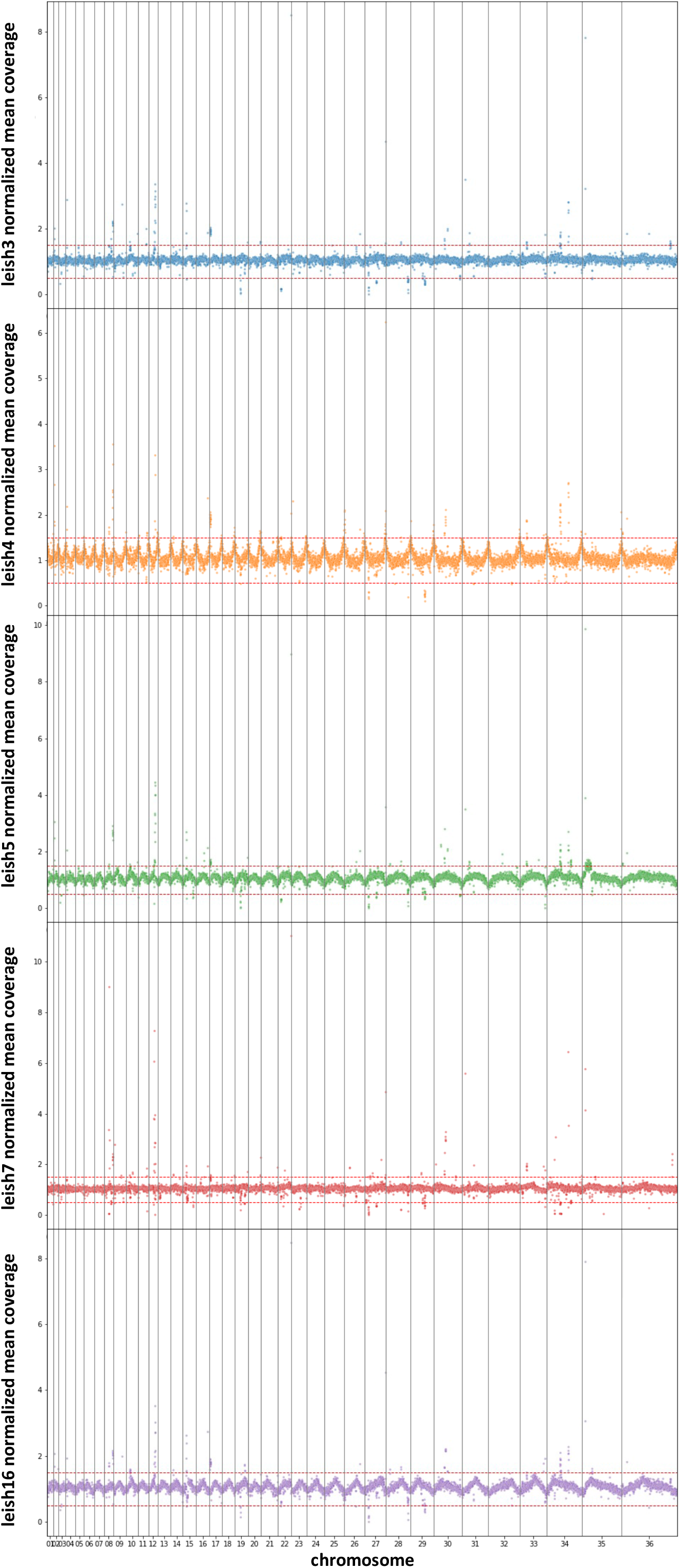
Scatterplots showing gene CNVs for the individual strains revealed by differences in normalized mean coverage. The X-axis represent genomic indices (their rank when sorted by genomic coordinates of their start positions). The Y-axis represents the genes’ normalized mean coverage (see methods).

**Figure S2:**
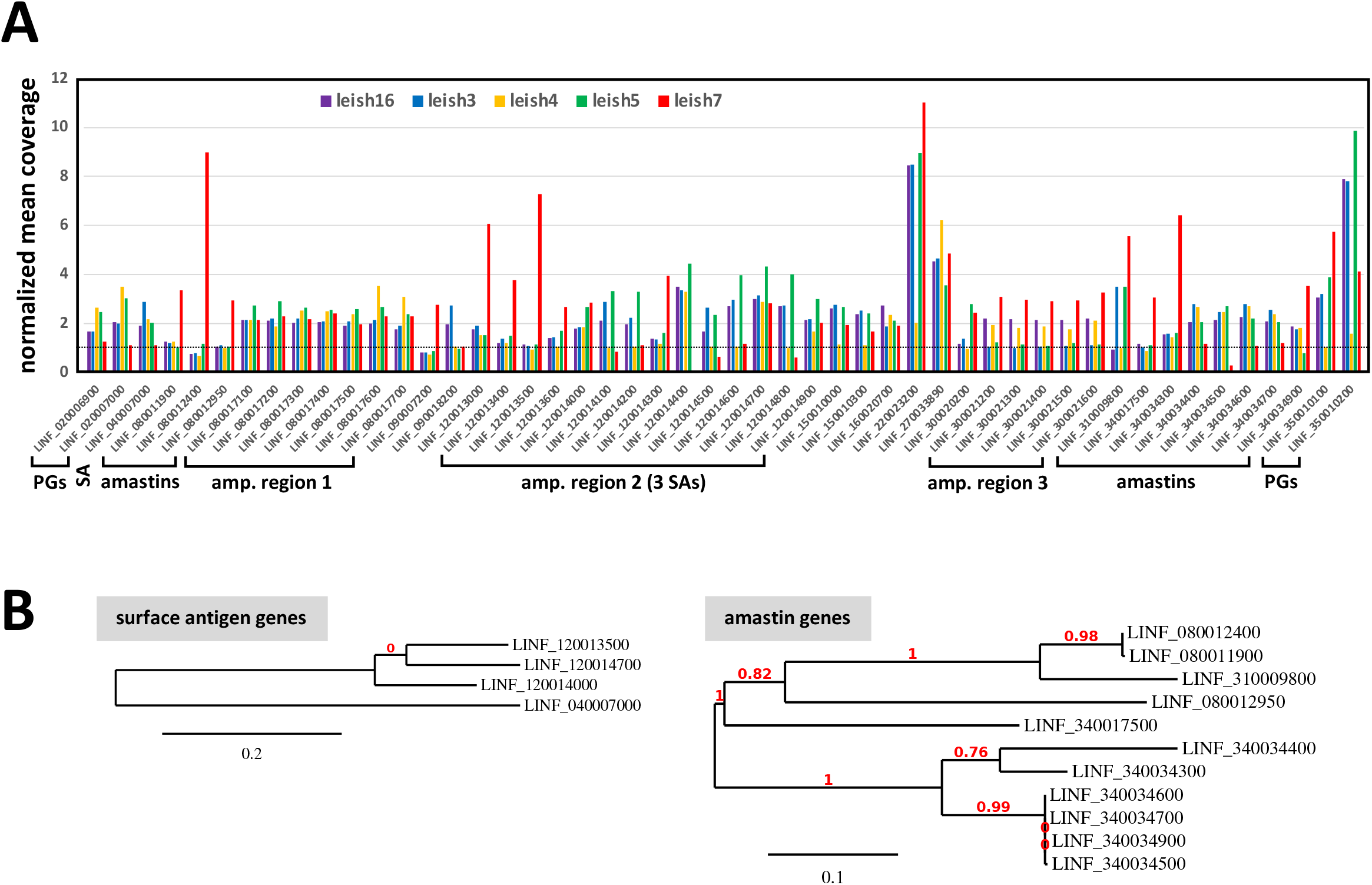
(A) Convergent amplifications revealed by similar coverage changes (Y-axis) for the genes indicated (X-axis) across the analysed strains (see colour code). (B) Phylogenetic analysis of the indicated genes performed on the www.phylogeny.fr web service using FASTA-formatted sequence files.

**Figure S3:**
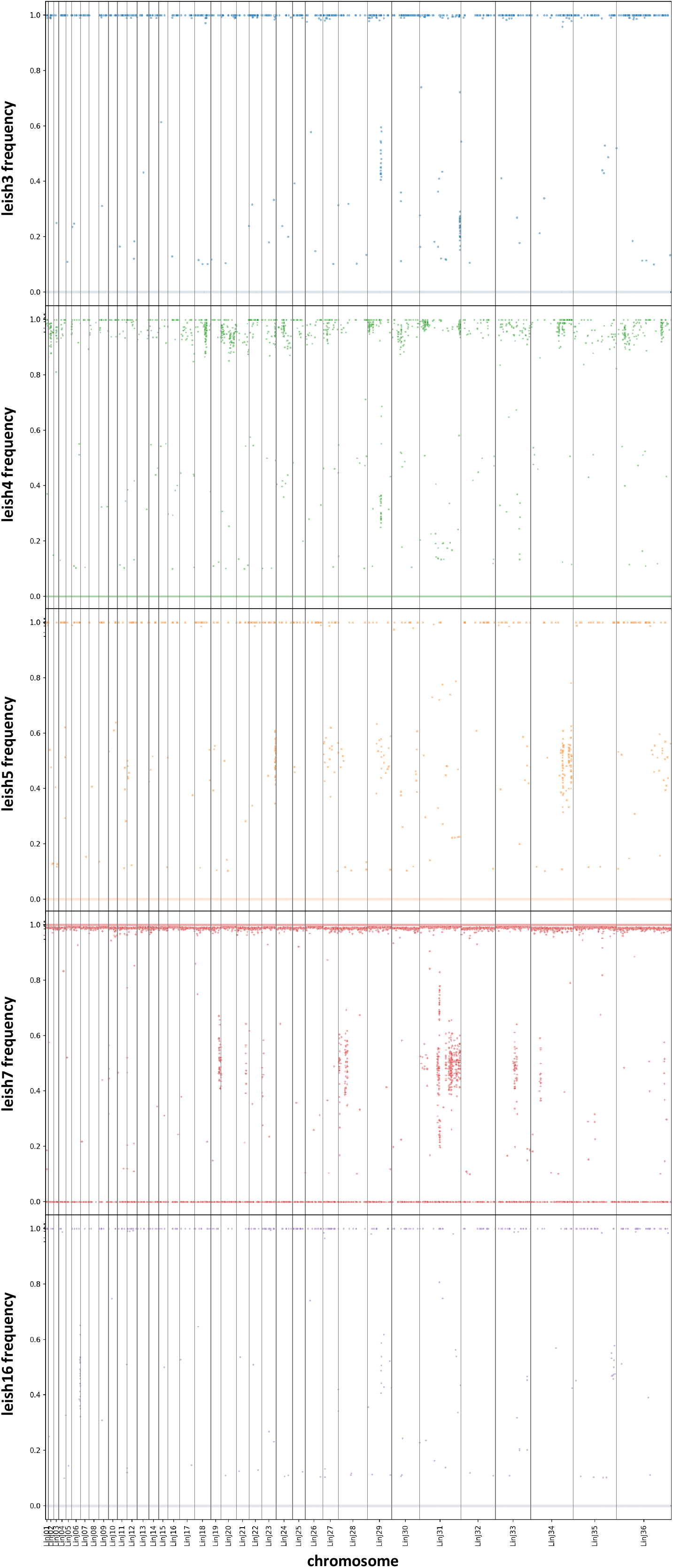
SNP scatterplots for the indicated strains (see Y-axis labeling). The Y-axis represents the SNPs genomic index (rank when sorted by genomic position) and the X-axis represents their frequency.

## List of supplementary tables

**Table S1:** Mapping metrics

**Table S2:** All gene CNVs

**Table S3:** Most significant gene CNVs

**Table S4:** Gene depletions

**Table S5:** SNP frequencies

**Table S6:** Designation, characteristics and MLMT profiles of the *Leishmania* strains

**Table S7:** *Leishmania donovani* complex strains used in this study

