## Supplementary material for "Genomic and epidemiological evidence for the emergence of a putative *L. donovani/L. infantum* hybrid with unusual epidemiology in Northern Italy": Table S7: Leishmania donovani complex strains used in this study

| **Table S7: *Leishmania donovani* complex strains used in this study** | | | | | |
| --- | --- | --- | --- | --- | --- |
| **Species** | **Lab code** | **Country** | **WHO-code** | **Zymodeme** | **Reference** |
| *Leishmania infantum* | H18 | Portugal | MHOM/PT/2004/IMT363 | MON-24 | [1] |
|  | H27 | Portugal | MHOM/PT/1997/IMT227 | MON-29 | [1] |
|  | H28 | Portugal | MHOM/PT/1997/IMT228 | MON-29 | [1] |
|  | H30 | Portugal | MHOM/PT/1992/IMT181 | MON-24 | [1] |
|  | H32 | Portugal | MHOM/PT/1989/IMT163 | MON-1 | [1] |
|  | H37 | Portugal | MHOM/PT/1993/IMT184 | MON-1 | [1] |
|  | H58 | Portugal | MHOM/PT/2004/IMT369 | MON-24 | [1] |
|  | C1 | Portugal | MCAN/PT/2003/IMT300 | MON-1 | [1] |
|  | C18 | Portugal | MCAN/PT/1995/IMT205 | MON-1 | [1] |
|  | C33 | Portugal | MCAN/PT/2003/IMT322 | MON-1 | [1] |
|  | FL4 | Portugal | IARI/PT/1989/IMT171 | MON-24 | [1] |
|  | FL5 | Portugal | IARI/PT/1989/IMT172 | MON-24 | [1] |
|  | ESP1 | Spain | MHOM/ES/93/PM1 | MON-1 | [1] |
|  | ESP2 | Spain | MHOM/ES/86/BCN16 | MON-1 | [1] |
|  | ESP3 | Spain | MHOM/ES/2001/LLM-981 | MON-1 | [1] |
|  | ESP4 | Spain | MCAN/ES/2001/LLM-1007 | MON-1 | [1] |
|  | ESP5 | Spain | MCAN/ES/2002/LLM-1139 | MON-1 | [1] |
|  | FR1 | France | MHOM/FR/97/LSL29 | MON-1 | [1] |
|  | FR2 | France | MHOM/FR/78/LEM75 | MON-1 | [1] |
|  | FR3 | France | MHOM/FR/80/LEM189 | MON-11 | [1] |
|  | GR1 | Greece | MCAN/GR/2002/GR9 | MON-1 | [1] |
|  | GR2 | Greece | MCAN/GR/2003/GR12 | MON-1 | [1] |
|  | GR3 | Greece | MHOM/GR/2003/GR19 | MON-98 | [1] |
|  | GR4 | Greece | MHOM/GR/2004/GR17 | MON-98 | [1] |
|  | GR5 | Greece | MHOM/GR/2002/GR26 | MON-1 | [1] |
|  | TUS227 | Tunisia | MHOM/TN/2002/Tus227 | MON-24 | [2] |
|  | PLV11 | Tunisia | MHOM/TN/2005/PLV11 | MON-24 | [2] |
|  | PLV15 | Tunisia | MHOM/TN/2005/PLV15 | MON-24 | [2] |
|  | PLV28 | Tunisia | MHOM/TN/2005/PLV28 | MON-24 | [2] |
|  | TUS167 | Tunisia | MHOM/TN/2001/Tus167 | MON-1 | [2] |
|  | 27S | Tunisia | MHOM/TN/2002/27M | MON-1 | [2] |
|  | 246M | Tunisia | MHOM/TN/2002/246M | MON-1 | [2] |
|  | 20S | Tunisia | MHOM/TN/2002/20S | MON-1 | [2] |
|  | 22MO | Tunisia | MHOM/TN/2002/22MO | MON-1 | [2] |
|  | TUS221 | Tunisia | MHOM/TN/2002/Tus221 | MON-1 | [2] |
|  | TUM222 | Tunisia | MHOM/TN/2002/Tum222 | MON-1 | [2] |
|  | LCnJ20S | Tunisia | MCAN/TN/2002/LCnJ20S | MON-1 | [2] |
|  | GGCH1 | Tunisia | MCAN/TN/2002/GGCH1/02 | MON-1 | [2] |
|  | LCnJ20G | Tunisia | MCAN/TN/2002/LCnJ20G | MON-1 | [2] |
|  | LC78 | Tunisia | MHOM/TN/2004/LC78 | MON-1 | [2] |
| *Leishmania donovani* | DON39 | India | MHOM/IN/0000/DEVI | MON-2 | [3] |
|  | DON1 | India | MHOM/IN/1980/DD8 | MON-2 | [3] |
|  | DON104 | Nepal | MHOM/NE/2003/BPK294/0 | nd | [3] |
|  | NLB189 | Kenia | MHOM/KE/1983/NLB189 | MON-37 | [4] |
|  | NLB323 | Kenia | MHOM/KE/1985/NLB323 | MON-37 | [4] |
|  | ET1 | Etiopia | MHOM/ET/72/GEBRE 1 | MON-82 | [1] |
|  | ET3 | Etiopia | MHOM/ET/67/LV9 | MON-18 | [1] |
|  | SD1 | Sudan | MHOM/SD/82/GILANI | MON-30 | [1] |
|  | SD2 | Sudan | MHOM/SD/62/3S | MON-81 | [1] |
|  | CD44cl.1 | Cyprus | MCAN/CY/2005/CD44cl.1 | MON37 | [5] |
|  | CH35 | Cyprus | MHOM/CY/2006/CH35 | MON37 | [5] |

***Footnote:*** *nd, not determined; [1] Cortes S et al, Infect Genet Evol 2014, 26: 20–31, PMID: 24815728; [2] Chargui N et al, International Journal for Parasitology 2009, 39: 801–811, PMID: 19211023; [3] Alam MZ et al, Infect Genet Evol 2009, 9(1): 24–31, PMID: 18957333; [4] Alam MZ et al, Microbes Infect. 2009, 11(6-7): 707–715, PMID: 19376262 ; [5] Gouzelou E et al, Parasit Vectors. 2013; 6:342, PMID: 24308691.*
